# GenOtoScope: Towards automating ACMG classification of variants associated with congenital hearing loss

**DOI:** 10.1101/2021.12.23.474074

**Authors:** Damianos P. Melidis, Christian Landgraf, Gunnar Schmidt, Anja Schöner-Heinisch, Sandra von Hardenberg, Anke Lesinski-Schiedat, Wolfgang Nejdl, Bernd Auber

## Abstract

Since next-generation sequencing (NGS) has become widely available, large gene panels containing up to several hundred genes can be sequenced cost-efficiently. However, the interpretation of the often large numbers of sequence variants detected when using NGS is laborious, prone to errors and often not comparable across laboratories. To overcome this challenge, the American College of Medical Genetics and Genomics and the Association for Molecular Pathology (ACMG/AMP) introduced standards and guidelines for the interpretation of sequencing variants. Further gene- and disease-specific refinements regarding hereditary hearing loss have been developed since then. With more than 200 genes associated with hearing disorders, the manual inspection of possible causative variants is especially difficult and time consuming. We developed an open-source bioinformatics tool `GenOtoScope`, which automates all ACMG/AMP criteria that can be assessed without further individual patient information or human curator investigation, including the refined loss of function criterion (“PVS1”). Two types of interfaces are provided: (i) a command line application to classify sequence variants in batches for a set of patients and (ii) a user-friendly website to classify single variants. We compared the performance of our tool with two other variant classification tools using two hearing loss data sets, which were manually annotated either by the ClinGen Hearing Loss Gene Curation Expert Panel or the diagnostics unit of our human genetics department. `GenOtoScope` achieved the best average accuracy and precision for both data sets. Compared to the second-best tool, `GenOtoScope` improved accuracy metric by 25.75% and 4.57% and precision metric by 52.11% and 12.13% on the two data sets respectively. The web interface is freely accessible. The command line application along with all source code, documentation and example outputs can be found via the project GitHub page.

**Author summary:** New high-throughput sequencing technologies can produce massive amounts of information and are utilized by laboratories to explain the often complex genetic aetiology of hereditary diseases. The most common sensory disease, hearing loss, is often hereditary and has a high impact on a patient’s every-day life. To use these sequencing technologies effectively, software tools were developed that can aid researchers interpreting genetic data by semi-automatically classifying the biologic (and thus potentially medical) impact of detected variants (the alterations of the patient’s genome compared to the human reference genome). The available genetic variant classification tools are either not designed specifically for the interpretation of variants detected in subjects with hearing loss or they do not allow researchers to use them for batch classification of all variants detected, e. g. in a study group. To address this drawback, we developed `GenOtoScope`, an open-source tool that automates the pathogenicity classification of variants potentially associated with congenital hearing loss. `GenOtoScope` can be applied for the automatic classification of all variants detected in a set of probands.

## Introduction

Due to the establishment of modern high-throughput next generation sequencing (NGS) technologies, an ever-increasing amount of sequencing data can be generated. Nevertheless, a whole exome sequencing (WES) file contains approximately 60,000 variants per proband. Consequently, laboratories have to overcome the hurdle of processing this vast amount of data to link the genotype to phenotype [1]. Notably, the manual classification of variants, by expert curators, is not only time-consuming, but even more, prone to inconsistent functional interpretation and pathogenicity classification of a variant between distinct laboratories [2].

To address this challenge, the American College of Medical Genetics and the Association for Molecular Pathology (ACMG/AMP) published a set of evidence-based criteria to classify patients variants in five classes of pathogenicity, “benign” (class 1), “likely benign” (class 2), “variants of uncertain significance” (“VUS”) (class 3), “likely pathogenic” (class 4), and “pathogenic” (class 5) [10]. To specialize for a diverse set of phenotypes with distinct penetrance, allelic and genetic heterogeneity, ACMG updated its classification criteria for specific hereditary diseases, for example hereditary (breast/ovarian) cancer [3] or cardiomyopathy [4], through the ClinGen Variant Curation Expert Panels (VCEP).

Hearing loss (HL) is the most common sensory disorder with a high impact on the quality of social and work life of the patient. A genetic aetiology can be linked to approximately 50% of the affected individuals [5]. Besides various forms of nonsyndromic hearing loss (NSHL) affecting only the function of the ear, HL can also be a symptom of a superordinate disorder involving other organ systems (syndromic hearing loss). Thus, HL is very heterogeneous with well over 100 genes known to be associated with monogenetic NSHL and more than 400 distinctive syndromes comprising HL as one of their characteristic symptoms as well [5].

To facilitate the challenging classification of variants for HL, [11] have published disease-specific evidence-based ACMG-criteria. Application of these adjusted criteria has been shown to achieve better classification performance compared to the standard evidence-based criteria for known HL-related variants [6]. A recently published bioinformatics tool, VIP-HL [8], automates 13 out of the 24 evidence-based criteria specified for HL. However, VIP-HL is an online tool that accepts only a single variant per time, thus hindering the automatic and time-efficient interpretation of all variants of WES files for a set of investigated patients, for a heterogeneous condition as HL.

To address this limitation of VIP-HL, we present `GenOtoScope`, a bioinformatic tool which accepts as input a genomic variant file (VCF) and computes the pathogenicity class and pathogenicity probability for each input variant, based on [11] and [22]. To this end, we designed and implemented algorithms to automate all the evidence-based criteria that need no further individual patient information or human curator investigation. This results to 12 implemented criteria, out of the 24 criteria in total, namely PVS1 (all strengths), PS1, PM1, PM2 (PM2 supporting), PM4, PM5 (PM5 strong), PP3, BA1, BS1 (BS1 supporting), BP3, BP4 and BP7. We will provide `GenOtoScope` as an open-source project, accessible as a command line application to classify the WES patients files and as an online tool to classify a single genomic variant of interest.

We benchmarked the performance of `GenOtoScope` compared to two established classification tools, InterVar [7] and VIP-HL, in two HL data sets. These data sets consist of manually curated HL variants. `GenOtoScope` outperformed the other two classification algorithms, both, in terms of accuracy and precision. Finally, we investigated the reasons for this best performance of `GenOtoScope`, by calculating the difference between the activation frequencies of a tool over the manual curation, for each evidence-based criterion.

In summary, our contributions are:

- Introduce `GenOtoScope` in two programming interfaces, a command line application for bioinformatics experts to classify WES VCF files of a set of patients and as web-based application for non-bioinformatics experts to classify single variants.
- Compare `GenOtoScope` classification performance to InterVar and VIP-HL for two manual annotated HL data sets.
- Make `GenOtoScope` an open-source bioinformatics tool, therefore enabling the research community to extend the tool for other diseases.

## Materials and methods

### Automating the examination of ACMG evidence criteria

`GenOtoScope` currently implements 12 out of 24 ACMG evidence-based criteria specified for hearing loss [11]. More specifically these criteria are PVS1 (all strengths), PS1, PM1, PM2 (PM2 supporting), PM4, PM5 (PM5 strong), PP3, BA1, BS1 (BS1 supporting), BP3, BP4 and BP7. Based on class category the implemented criteria are sorted in 7 pathogenic and 5 benign criteria. With respect to the data types needed for ACMG criteria, we categorize our implemented criteria into 3 population data criteria, 8 computational and predictive data criteria and 1 functional data criterion.

The unimplemented criteria by `GenOtoScope` are 12. These criteria are: PS2, PS3, PS4, PM3, PM6, PP1, PP4, BS2, BS3, BS4, BP2 and BP5. The main reasons not to implement these criteria are: (i) the lack of established processing algorithm (ii) the lack of data and (ii) further patient information. That is, for the criteria needing functional data, PS3 and BS3, there are no established algorithms that can automatically extract the result of a functional study publication, for a given human variant. As the lack of data is concerned, the examination of the PS4 criterion cannot be automated as there is no database to contain the prevalence of affected and control individuals for all possible variant types. Equally, there is no database with the respective information to automate BS2 and BP2 criteria. Last, the need for genomic data from the patient’s family disables the examination of the segregation data criteria: PS2, PM3, PM6, PP1, PP4, BS4 and BP5.

The number of missing implemented criteria is competitive with the other classifications tools. VIP-HL implements only one extra criterion, BS2. The main reason not to also implement BS2 is that VIP-HL uses particular thresholds, which are not specified by ACMG HL original work, and thus it may not reflect all penetrance and inheritance modes of all HL-related genes.

InterVar implements 18 out of the 24 ACMG original criteria [10]. As explained in the introduction, disease-specific ACMG criteria may vary from the original 24. Therefore the PP2, PP5, BP1 and BP6 criteria automated by InterVar are not applicable for HL. The remaining two criteria automated by InterVar and not by `GenOtoScope` are PS4 and BS2. To automate these criteria, InterVar used the ANNOVAR annotation tool [35]. However, this tool implements PS4 using a general threshold on a phenotype-based GWAS catalog, consequently the called enriched pathogenic variants may not include all HL-relevant variants. Similarly, to automate BS2 criterion, InterVar uses the zygotic information of a healthy individual in the 1000 Genomes project [36] based on the inheritance mode of the variant. Nevertheless, specific thresholds of healthy individuals should be used for HL, which are not published by [11]. As a consequence, there may be false negative cases; InterVar should activate PS4 or BS2 for a given HL variant, but it may not. Finally, the remaining criteria need manual curation or additional information not publicly available (e.g. segregation or phenotypic data), therefore they are not implemented by any of the three classification tools.

In our thorough evaluation, shown in the results section, we demonstrate that regarding the 12 ACMG criteria processed by either tool, `GenOtoScope` achieved the best averaged accuracy and precision scores for both tested data sets. This is due to the activation frequency of these criteria being much closer to human curation in `GenOtoScope` than in VIP-HL and InterVar, which trigger the commonly implemented 12 criteria much less frequently.

To sum up, our choice to implement these 12 criteria, which are refined for HL, can lead to standardized classification results for all HL-relevant genes. Besides, our implementation of the criteria presents two more advantages: In contrast to the usage of the ANNOVAR annotation tool, licensed for commercial use, we construct all annotation files needed to examine the ACMG criteria, using freely accessible databases and offer `GenOtoScope` with an open-source software license. Therefore, any interested researcher can update the corresponding code section to produce adjusted annotations to her needs. Equally, the researcher can update the code to change the steps used to examine a given criterion. The second advantage is that `GenOtoScope` (like VIP-HL) outputs comments for each examined criterion, whereas InterVar does not. This extra information can facilitate the variant curator to justify the activation of a criterion and thus increace the explainability of the classification.

### GenOtoScope workflow

In the following, the methodology to implement the ACMG evidence-based criteria for congenital hearing loss is explained in five key steps. The conceptual workflow of the web and command line interface (CLI) of `GenOtoScope` is depicted in Fig 1.

**Fig 1.**
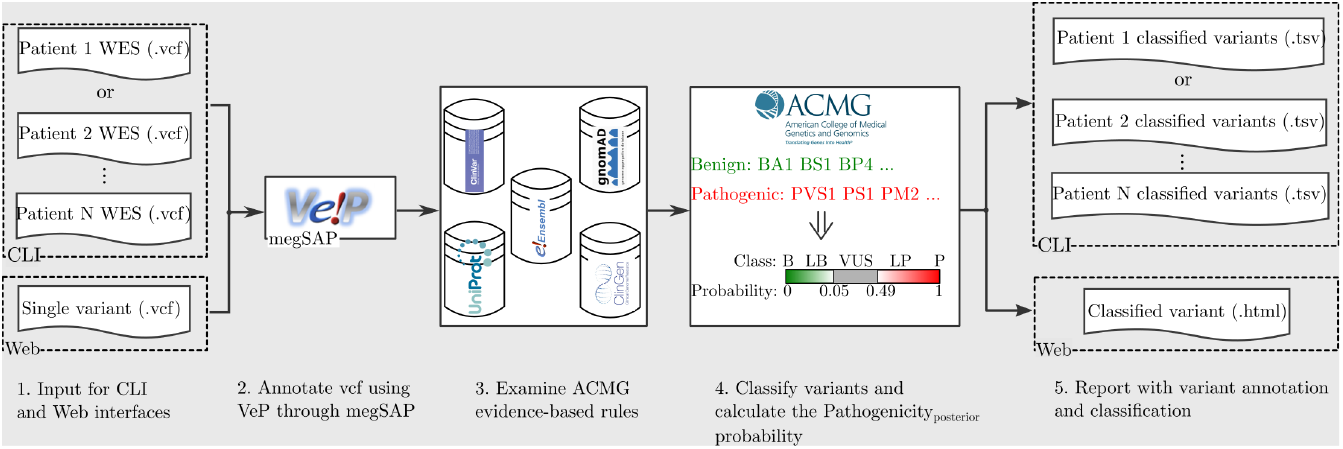
Conceptual workflow of GenOtoScope.

In the first step, the user inputs a variant file (VCF), which, depending on the used interface, may contain a single variant or a larger set of variants of a patient (e.g. full WES data set). Multiple VCFs can be submitted simultaneously.

Next, functional annotation of the VCF takes place using the VeP annotation tool [33] through the megSAP bioinformatics application^1^. The resulted intermediate variant file is organized as a standard matrix file (tabular file) where each row is a variant and a column contains variant annotation. These columns contain the basic variant information (for example chromosome position of variant and affected gene name), the transcript and the protein HGVS signature and the unique functional annotation for the variant (e.g. minor allele frequency in gnomAD subpopulations [20], the OMIM variant observed clinical description [34] and the REVEL pathogenicity score for the variant [15]).

The third step uses the core sub-algorithms of `GenOtoScope` to automatically analyze the listed variants according to ACMG criteria. These sub-algorithms access programmatically four databases: the human clinical variants database ClinVar [37], the human exomes database gnomAD [20], the protein knowledge database UniProt [38] and the clinical genome database [39]. Extracted annotations are organized based on the Ensembl features [21] for a variant-affected transcript. Beyond the mere result of checking a criterion (activation or non-activation), the tool stores a descriptive comment on the reason for activation or non-activation, to be used as an explanation for the user.

In the following step, the tool combines the activated evidence-based criteria to classify the variant into 5 pathogenicity categories (“benign”, “likely benign”, “VUS”, “likely pathogenic” and “pathogenic”) according to ACMG guidelines. If none of the criteria is activated, the tool classifies the variant as VUS. Subsequently, in the same fourth step, `GenOtoScope` computes the pathogenicity posterior probability based on [22]. This is intended to allow a better discrimination of VUS and additional re-classification of VUS into benign or pathogenic variants.

In the fifth and final step, `GenOtoScope` extends the intermediate annotation tabular file with the criteria activation results and the comments along with the predicted ACMG class and the computed pathogenicity probability. Finally, the tool will save this file as the produced classification output.

A crucial sub-step of this workflow is the construction of annotation files, which is needed for the automatization of the examination of the ACMG evidence-based criteria. We constructed the needed annotation files, clinical-significant exons, HL-relevant transcripts, critical regions for proteins, critical regions for proteins without benign variants and protein repeat regions without domain intersection, using publicly available data sets.

### GenOtoScope interfaces

Having described the general workflow, we continue with the presentation of the two interfaces.

#### Web interface

The web application is targeted for free online usage. Advanced bioinformatics skills are not required. Screenshots of the home page and of an example output page, of the `GenOtoScope` website, are shown in Fig 2. At the home page, users can upload a single variant file (VCF). The website will annotate and convert the VCF to GSvar file through the megSAP application. A result page (.html) will be generated to show: (i) the variant’s ACMG classification and computed pathogenicity posterior probability for all known inheritance modes of the affected gene, (ii) the list of all examined ACMG criteria along with comments on their activation or deactivation and (iii) the basic information for variant and affected transcripts.

**Fig 2.**
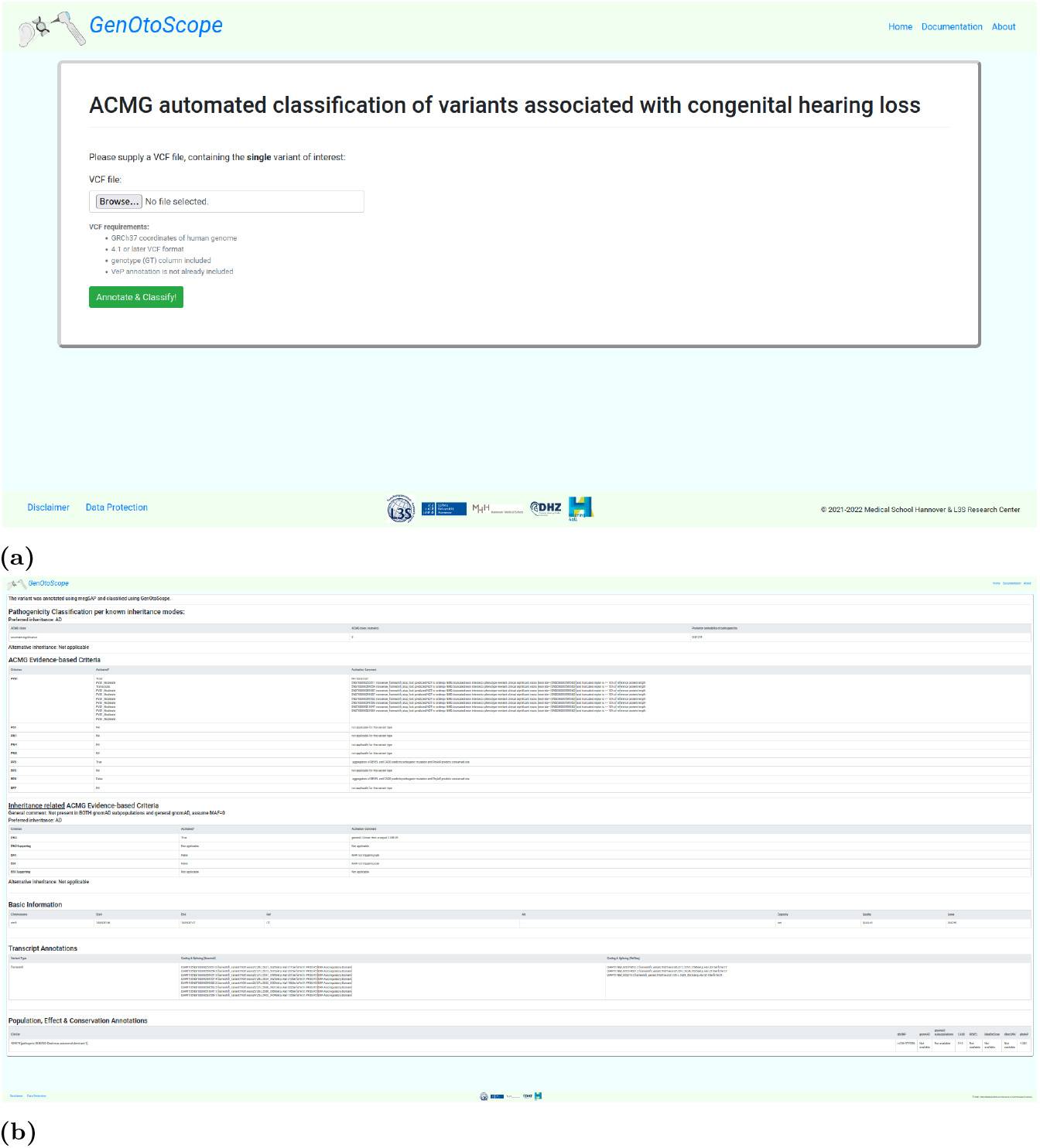
Web interface of GenOtoScope. (a): The home page of the `GenOtoScope` website. (b): The output page, for an example variant (RS id: 1064797096), which includes its classification based on HL-specified ACMG guidelines.

#### Command line interface

The command line interface (CLI) is tailored to bioinformatics personnel. The first command of this mode, `genotoscope annotate.py`, will accept as input a folder of VCF files or a single VCF file. It will annotate the input VCF files and convert them into GSvar files. The second command, `genotoscope classify.py`, will accept as input a folder of GSvar files or a single GSvar file, the output of the previous command. Then it will automatically examine the ACMG evidence-based criteria to classify and compute the pathogenicity posterior probability for each variant in an input GSvar file. The output will be an extended GSvar file containing information on the examination of the ACMG evidence-based criteria, the ACMG pathogenicity class and the pathogenicity posterior probability. Running examples of these two commands are shown in Fig 3.

**Fig 3.**
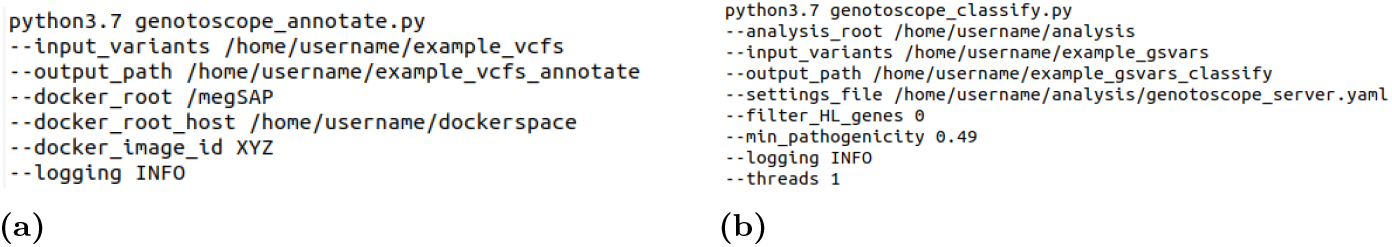
Command line examples for the two commands of GenOtoScope. (a) Annotate all variants presented in VCF files, in input folder, using megSAP application and save results in GSvar files. (b) Classify all variants presented in GSvar files based on ACMG guidelines specified for HL.

#### Automating examination of ACMG evidence-based criteria

In the following subsections, we briefly describe our implementation of the aforementioned 12 ACMG criteria: PVS1 is automated based on [13]. Information from ClinVar database is used for the implementation of PS1 and PM5 (including PM5 Strong). Automation of PM1 examines critical regions provided by [11] and a purpose-built annotation file containing critical regions without benign mutations. Customized annotation files are also used for (non) repetitive region dependent criteria PM4 and BP3, whereas automation of PP3, BP4 and BP7 employs established prediction algorithms. Population frequency data for implementation of PM2 (PM2 Supporting), BA1 and BS1 (BS1 Supporting) is taken from gnomAD database.

#### Refined PVS1

PVS1 criterion is assessed for start-loss, nonsense (stop gained), stop-loss, frameshift, in-frame, splice acceptor and donor variants according to [13].

First, the occurrence of nonsense-mediated decay (NMD) is predicted by a subroutine for each affected transcript using the HGVS signature of the variant to create the observed coding sequence per exon. Altered region is defined as variant-affected coding region. The algorithm locates the 5’-closest stop codon and follows the scheme published by [14] to assess impact of this premature termination codon (PTC) on NMD: Observed coding sequence is rated to escape NMD if PTC appears either within the 50 last bases of the penultimate exon or at most 200 bases downstream from the start codon or the transcript contains no introns. Otherwise, NMD is classified to occur. Fig. 4 illustrates this subprocess.

**Fig 4.**
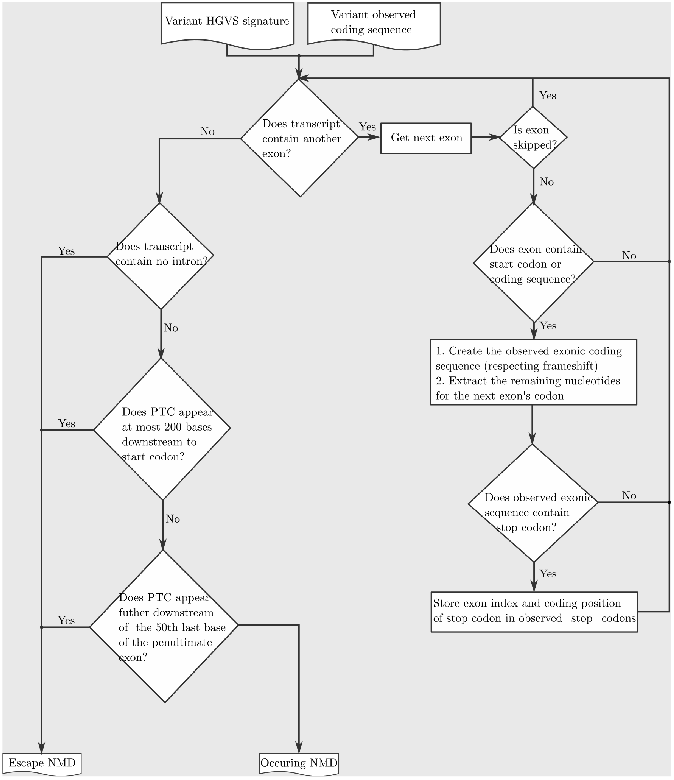
Conceptual flowchart to assess NMD for the refined PVS1 rule.

If NMD is predicted to occur, the algorithm intersects the stored variant-affected coding region to phenotype-relevant transcripts to decide the PVS1 outcome. If NMD is not predicted to occur, it intersects the variant-affected coding region with protein domain regions. If there is an intersection, it examines if the affected region overlaps a critical domain for protein function, to decide the PVS1 outcome. If the affected region is not within a known domain, overlap with clinically significant exons and phenotype-relevant transcripts is examined. If this is confirmed, it is investigated whether the PTC results in the removal of more than 10% of the reference protein product.

For start-loss variants, the algorithm first checks if any other transcript contains an alternative start codon. If not, it extracts potential in-frame start codons that are no further than 200 bases downstream of the lost start codon. Next, it queries ClinVar for pathogenic entries with at least one review star between the lost start codon and the detected in-frame start codon. If there is such a ClinVar entry, PVS1 (Moderate) is triggered, otherwise PVS1 (Supporting) is activated.

See supplementary for annotation files (clinically significant exons, phenotype-relevant transcripts and clinically significant exons).

#### PS1 and PM5 (PM5 Strong)

The workflow of assessing the PS1 and PM5 criteria is shown in Fig. 5.

**Fig 5.**
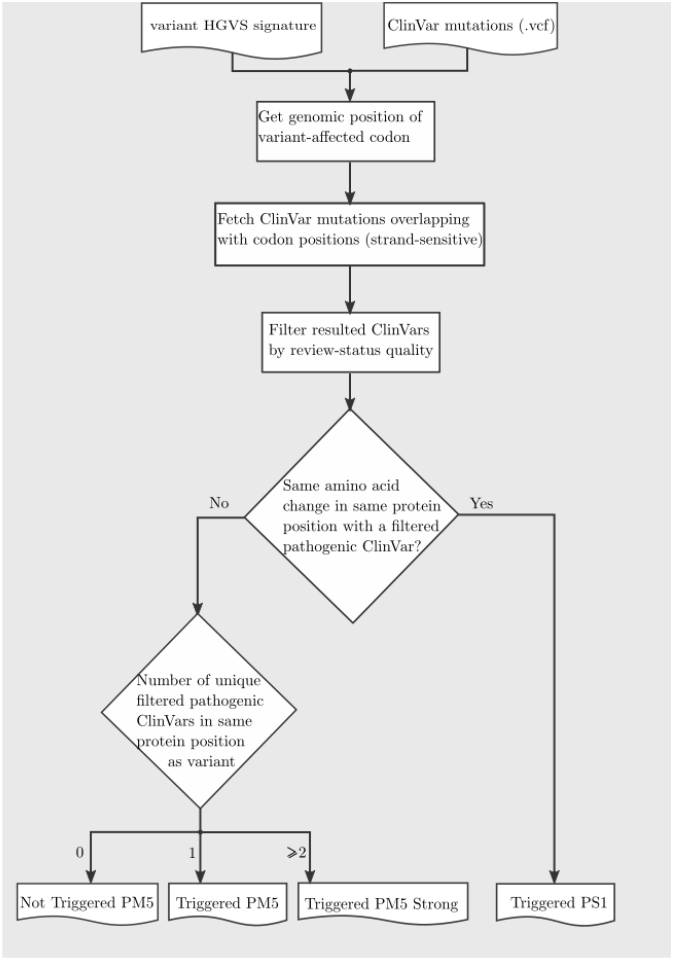
Conceptual flowchart for examining PS1 and PM5 (PM5 Strong).

First, genomic positions of the affected codon are computed based on exonic variant location and directionality of the respective gene. Then, all missense variants at corresponding genomic positions are extracted from ClinVar and filtered by strand to match the directionality of the affected gene. Additionally, ClinVar entries can be filtered by review status as the user can define a minimum number of quality stars as a threshold for variants to be considered (default value: 1).

The filtered variants and resulting amino acids are used for further assessment of PS1 and PM5 criteria: PS1 is triggered if any filtered-in variant from ClinVar that is rated as pathogenic results in the same amino acid change as the observed variant. PM5 is triggered if the filtered-in ClinVar entries do not contain the observed amino acid change but at least one pathogenic variant affecting the same codon. If the entries include two or more such variants, PM5 is applied as strong evidence (PM5 Strong) according to [11].

#### PM1

For automation of PM1 a custom-made annotation file is used. It comprises all critical protein regions without benign ClinVar entries and also includes specific domains and motifs of hearing loss related proteins as defined by [11]. Please see supplementary for detailed description of used hotspot reagions related to HL.

#### PM4

The algorithm for this criterion is applied to all in-frame (deletions/duplications) and stop-loss variants that do not trigger PVS1 in any strength level. Considering PTC assessed in PVS1 subroutine, length of observed proteins is calculated and compared to reference protein length. For length differences greater than 10%, PM4 is triggered except for variants in known repetitive regions derived from UniProt.

#### BP3

The algorithm for this criterion is applied to all in-frame (deletions/duplications) and nonsense (stop gained) variants. Based on an annotation file containing functional domains and repeat regions derived from UniProt, BM3 is triggered if the variant-affected coding region overlaps a repetitive region without known function.

#### PP3, BP4 and BP7

`GenOtoScope` incorporates in silico tools for conservation (PhyloP [17]), splicing (MaxEntScan [18], dbscSNV [19]) and missense-prediction (REVEL [15], CADD [16]). See supplementary for aggregation of scores and thresholds.

For missense variants, activation of PP3 requires positive pathogenicity prediction (REVEL/CADD) and high conservation (PhyloP) scores. In contrast, BP4 is triggered if conservation and predicted probability of pathogenicity are low.

Variants with no immediate impact on amino acid sequence (exclusion: canonical splice site variants) are similarly screened for potential effects on splicing. If splicing is predicted to be affected and the nucleotide is highly conserved, PP3 is activated. Conversely, if a potential splice variant is predicted to have no splicing effect and conservation is low, BP7 is triggered for synonymous variants and BP4 for other variant types respectively.

#### BP7

The BP7 rule is evaluated for all synonymous variants following the similar procedure as for PP3 and BP4. First, the algorithm examines the splicing effect based on majority-voting of the predictions by the MaxEntScan or the dbscSNV tools. Then, it determines the pathogenicity of the variant based on majority-voting on the Revel or CADD predictions. Finally, it examines the conservation of the variant site by PhyloP score. The BP7 rule is triggered if the variant is predicted to have no splicing effect, to be not pathogenic and is not highly conserved, otherwise BP7 is not triggered.

#### PM2 (PM2 Supporting) BA1 and BS1 (BS1 Supporting)

Assessment of population data criteria uses adjustable minor allele frequency (MAF) thresholds, which by default are the ones defined by [11]. Each gene can be assigned a preferred mode of inheritance, which can be customized by providing an input file. Default settings comprise the inheritance modes of 164 hearing loss gene-disease pairs defined by the ClinGen Hearing Loss Gene Curation Expert Panel [24] plus preferred inheritance patterns for additional genes specified by the HG department of MHH. We will refer to ClinGen Hearing Loss Gene Curation Expert Panel commitee as VCEP-HL for convenience.

For each variant, allele frequencies (AF) of gnomAD subpopulations are retrieved. Known pathogenic variants with high AF are excluded from further assessment of BA1 and BS1 according to [11]. AF of each subpopulation and the median AF of all subpopulations are evaluated with respect to the appropriate inheritance mode threshold. PM2 (PM2 Supporting), BA1 and BS1 (BS1 Supporting) are triggered, if any subpopulation’s AF or the median AF matches the respective inheritance mode threshold.

#### Hearing-loss specific ACMG classification

Having assessed all applicable criteria for a given genomic variant, `GenOtoScope` combines the activated criteria to compute the respective ACMG class using the five-tier terminology system (“benign”, “likely benign”, “VUS”, “likely pathogenic” and “pathogenic”) defined by [10].

Moreover, `GenOtoScope` incorporates the extended recommendations of VCEP-HL for the following criteria combinations: (i) Variants triggering PVS1 and PM2 (Supporting) will be classified as “likely pathogenic” for genes associated with autosomal recessive inheritance. (ii) Variants activating BS1 without triggering any pathogenic criterion will be classified as “likely benign”.

#### Computation of pathogenicity probability

This feature is particularly intended for variants classified as VUS, due to insufficient or conflicting triggered evidence criteria. To help human curators discriminate the potential pathogenicity of VUS in a quantitative manner, `GenOtoScope` calculates the pathogenicity probability for each variant following [22].

To do so, the tool applies the naive Bayes model to calculate the posterior probability of pathogenicity given the triggered ACMG evidence rules using the following equations:

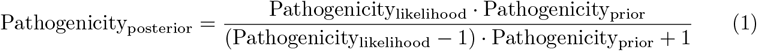

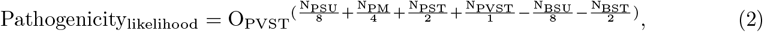

where the default parameters are used: Pathogenicity_prior_ = 0.1, O_PVST_ = 350 and X = 2.

The calculation of the pathogenicity probability is calculated automatically for all input variants.

## Results and Discussion

### Variant classification

#### Data sets

`GenOtoScope` variant classification was compared to similar tools: (1) InterVar, a tool for variant classification tested across a spectrum of phenotypes [7]; (2) VIP-HL, the recently published tool for hearing loss [8]. We benchmarked the accuracy and precision of variant classification on two data sets. The first data set is the publicly available set of manually annotated variants by ClinGen VCEP-HL [6], hereafter referred as VCEP-HL data set. This data set contains manual annotation for 158 variants associated with HL. These variants involve in 9 HL-relevant genes (*USH2A, COCH, GJB2, KCN☯4, MYO7A, MYO6, TECTA, SLC26A4* and *CDH23*). The second data set is the private set of manually annotated variants by the HG department of MHH, hereafter referred to as MHH data set. The MHH data set consists of 118 variants, which involve 36 HL-relevant genes. The included genes are: *COL11A1, USH2A, NLRP3, OTOF, ALMS1,PAX3, ILDR1, WFS1, COL11A2, COL9A1, MYO6, SLC26A4, CHD7, GRHL2, TMC1, WHRN, TNC, MYO3A, PCDH15, CDH23, OTOG, MYO7A, TECTA, COL2A1, MYO1A, P2RX2, GJB2, GJB6, ACTG1, MYH14, KCNE1, TMPRSS3, MYH9, SOX10, POU3F4* and *PRPS1*.

#### Performance metrics

To assess the prediction performance, we grouped “benign” and “likely benign” classes to “Benign”, “pathogenic” and “likely pathogenic” classes to “Pathogenic”. Thus, we created a three-class prediction task, containing the “Benign”, “Pathogenic” and “VUS” as possible classes.

Following the evaluation of the classification tool TAPES [26], we evaluate the accuracy of each software tool, using the area under the curve (AUC) metric of the Receiver Operating Characteristics (ROC) curve for all possible, one versus rest (of classes), predictions. To compare the precision of each algorithm, we used the AUC metric of the precision-recall curve. To explain the observed differences in prediction performances, we plotted the frequency of triggered rules by each tool and the manual curation.

#### Refined classification of VUS

We acknowledge that not all evidence-based criteria for HL need a manual curation, therefore cannot be automated. To evaluate the `GenOtoScope` classification potential with the currently implemented half of all evidence-based criteria for HL, we provided a refined classification based on pathogenicity probability. We will refer to this extended version of `GenOtoScope` as `GenOtoScope pathogenicity probability` and as `GenOtoScope prob` for short. In this classifier version, we reclassified the “VUS” variants, classified by the `GenOtoScope` original version, by their calculated pathogenicity probability. That is, `GenOtoScope pathogenicity probability` maintains the `GenOtoScope` classification for “benign”, “likely benign”, “likely pathogenic”, “pathogenic” and refines the variants classified as “VUS” to “likely benign” if Pathogenicity_posterior_ *≤* 0.05072, to “likely pathogenic” if Pathogenicity_posterior_ *≥* 0.49988. For the remaining cases, 0.05072 *<* Pathogenicity_posterior_ *<* 0.49988, the `GenOtoScope pathogenicity probability` keeps the “VUS” classification unchanged.

We have chosen these threshold values based on relaxing the lowest combination of the triggered criteria for the broader classes of *Pathogenic* and *Benign*. We then transformed this relaxed combination of criteria to pathogenicity probability based on Eq. 2. That is, for the *Pathogenic* broader class, the combination of the criteria, with the least pathogenicity strength resulting in *likely pathogenic* class, is *1 pathogenic moderate criterion and at least 4 pathogenic supporting criteria*. Based on available open data and further patient genetic data we have implemented seven out of the fourteen applicable ACMG criteria. Thus, we lowered the combination to *1 Moderate and 1 Supporting criterion*, which translates to the probability of 0.49988. Therefore, the `GenOtoScope pathogenicity probability` will refine the *VUS* class, by the original GenOtoScope, to *Pathogenic* for a variant with pathogenicity probability greater than or equal to 0.49988.

Similarly, for the *Benign* broader class, the combination of criteria with the lowest strength is *at least two benign supporting criteria* and results in the *likely benign* class. For the same reasons, the `GenOtoScope` currently implements five out of the total ten applicable criteria. Therefore, we reduced the requirements of this combination to *one benign supporting criteria* which translates to the pathogenicity probability of 0.05072. Consequently, `GenOtoScope pathogenicity probability` will reclassify a variant classified as *VUS*, by `GenOtoScope` original version, to *Benign* broader class if the variant’s probability is lower or equal to 0.05072.

#### Investigation of performance discrepancies

We sought out to investigate the reasons for the discrepancy in prediction performance between the classification tools. To do so, we extended the troubleshooting plots of [27], by calculating the log ratio of the activation frequency of an evidence-criterion by a classification tool and the manual curation, as:

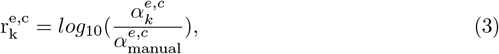

where 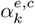 is the activation frequency of *e*, any of the implemented ACMG rules, by a tool *k* = {InterVar, VIP-HL, GenOtoScope} for a grouped class *c* ={pathogenic, VUS, benign}.

We computed all log ratios for each evidence rule, *e*, by each classification tool for the three grouped classes, *c*. Finally, we used heatmap plots to depict these log ratios.

### VCEP-HL data set

The ROC and precision-recall curves are shown in Fig 6. We observe that `GenOtoScope` and `GenOtoScope pathogenicity probability` achieved the best AUC scores for all three classes. In Precision-Recall curves, VIP-HL achieved slightly higher AUC compared to `GenOtoScope` for the “Benign” broader class. However, for the other two classes again `GenOtoScope` and `GenOtoScope pathogenicity probability` achieved the best AUC scores of the Precision-Recall curves.

**Fig 6.**
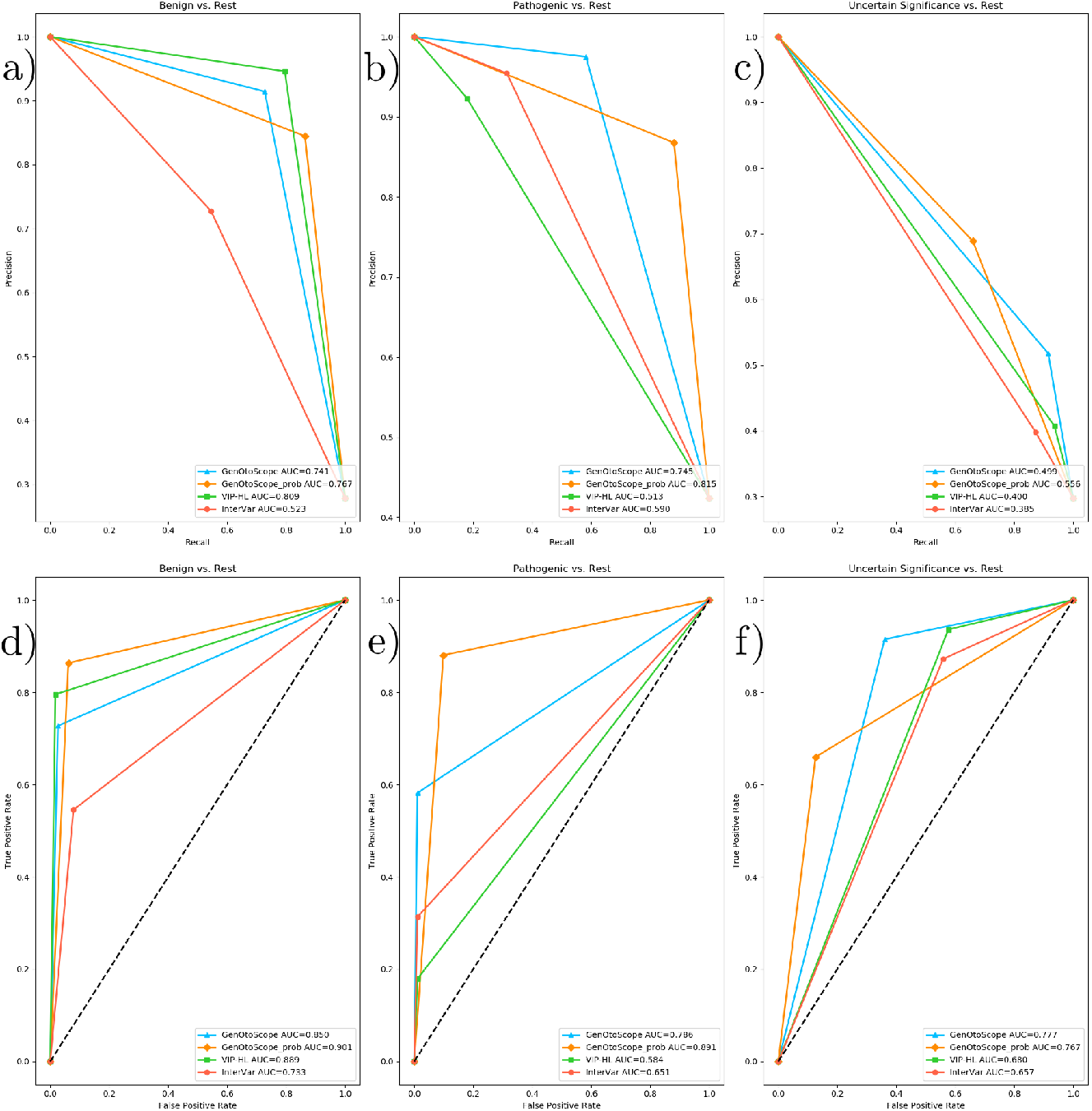
ROC curves and precision-recall curves for the VCEP-HL data set. a-c) ROC curve and AUC of all classification tools for VCEP-HL data set. (a) Prediction of the “Benign” broader class versus the “Pathogenic” broader class and the VUS class (b) Prediction of the “Pathogenic” broader class versus “Benign” broader class and the VUS class (c) Prediction the “VUS” class versus the “Benign” broader class and the “Pathogenic” broader class. d-f) Precision-recall curve and AUC of all classification tools for the VCEP-HL data set. (d) Prediction of “Benign” broader class versus the “Pathogenic” broader class and the VUS class (e) Prediction of the “Pathogenic” broader class versus “Benign” broader class and the VUS class (f)

Prediction of the “VUS” class versus the “Benign” broader class and the “Pathogenic” broader class. Besides, we calculated the performance scores, AUC of ROC and the average precision of the precision-recall curves for all classification tools. We show the micro-averaged scores, over the three broader classes (“Benign”, “VUS”, “Pathogenic”) in Table 1. Based on this table, the two versions of `GenOtoScope` classification achieved the best results for both AUC of ROC and the average precision.

**Table 1.**
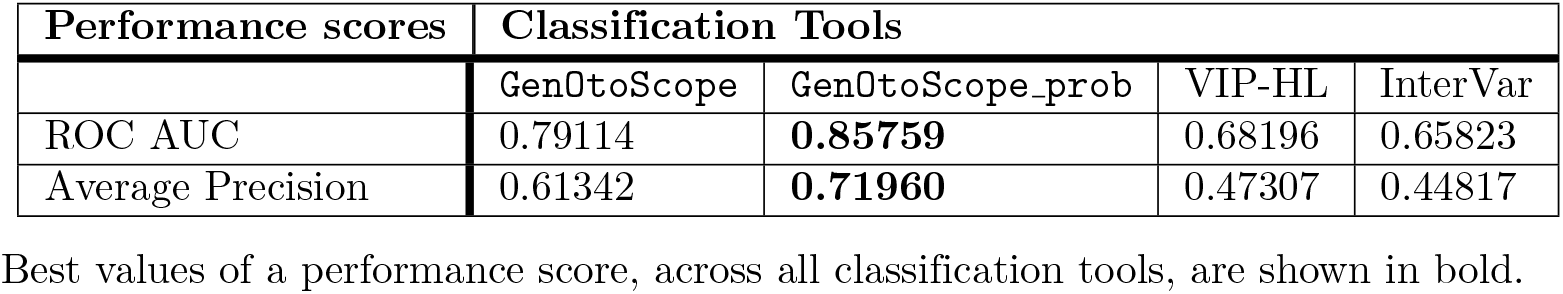
Micro-averaged performance scores for all classification tools, over the three broader classes in the VCEP-HL data set.

To explain the difference in prediction performance, we plot the heatmaps of the log ratio of activation frequency between a classification tool and the manual curation (3). The results are shown in Fig 7.

**Fig 7.**
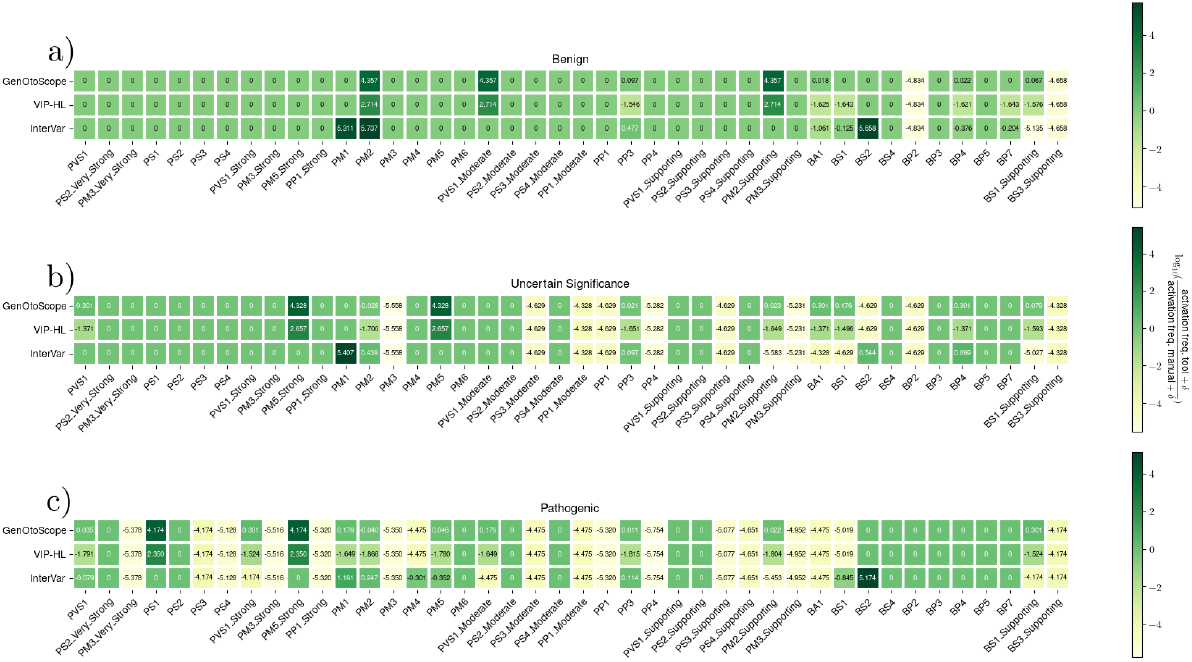
Activation frequency ratios for VCEP-HL data set. Log ratios calculated for each of the three classes classified by the VCEP-HL: (a) the “Benign” broader class, (b) the “VUS” class and (c) the “Pathogenic” broader class.

We observed the following patterns for each grouped class. First, for the “Pathogenic” broader class, VIP-HL activated 8 implemented pathogenic rules (PVS1 (Strong), PVS1 (Moderate), PM1, PM5, PVS1 and PM2) from 32 times less (PVS1 (Strong)) to 79 times less (PM2) than the manual curation. Nevertheless, `GenOtoScope` activated 5 out of these 8 rules with the same frequency as the manual curation (PVS1, PM2, PP3, PM2 (Supporting) and PM5). It activated the remaining 3 rules (PVS1 (Moderate), PM1 and PVS1 (Strong)) approximately twice as much as the manual curation.

For the “VUS” class, we observed that VIP-HL activated 8 implemented rules (BP4, BA1, PVS1, BS1, PM2 (Supporting), PP3 and PM2) from 25 times less (BP4) to 50 times less (PM2) than the manual curation. In contrast, `GenOtoScope` activated 3 out of the 8 rules (PM2, PP3, PM2 (Supporting)) with the same frequency as the manual curation and it activated the remaining 5 rules (BS1 (Supporting), BS1, BA1, BP4, PVS1) approximately one to two times more frequently than the manual curation.

For the “Benign” broader class, VIP-HL activated 6 implemented rules (PP3, BS1 (Supporting), BP7, BP4, BS1 and BA1) from 32 times less (BA1, BS1, BP4, BP7, BS1 (Supporting)) to 40 times less (PP3) than the manual curation. `GenOtoScope` activated 4 of these rules (BS1,BP7,BA1,BP4) with approximately the same frequency as the manual curation. The other two rules (PP3 and BS1 (Supporting)) were activated by genotoscope, one time more frequently than the manual annotation.

To examine the reasons for the lower precision of `GenOtoScope` for the “Benign” broader class compared to VIP-HL. We examined `GenOtoScope` misclassifications for this grouped class. That is, we examined the variants belonging in the VUS class and misclassified in the “Benign” broader class by `GenOtoScope`. These misclassified variants are seven, a significant amount on the calculation of precision for the total of 44 variants in the “Benign” broader class. The main reason for the misclassification was that manual annotation used not implemented rules to classify these variants as VUS. More specifically, for five out of the seven variants the manual curation used rules not implemented by `GenOtoScope` (for example PP1, PP4 or PM3) to classify the variants as VUS. The last two variants were misclassified by `GenOtoScope pathogenicity probability` as their calculated probability was lower than the set threshold for refining a variant classified from “VUS” into the “Benign” broader class. For completeness, VIP-HL classified correctly, as “VUS”, four out of these seven variants.

### MHH data set

The ROC curve and AUC scores are shown in Fig 8. In ROC curves, `GenOtoScope` or `GenOtoScope pathogenicity probability` scored the highest performance values, compared to VIP-HL and InterVar, for all three classes. In the Precision-Recall curves, `GenOtoScope` outperformed all classification tools, in terms of AUC score, for benign classification. `GenOtoScope` and `GenOtoScope pathogenicity probability` outperformed all classification tools, in AUC score for pathogenic and VUS classes.

**Fig 8.**
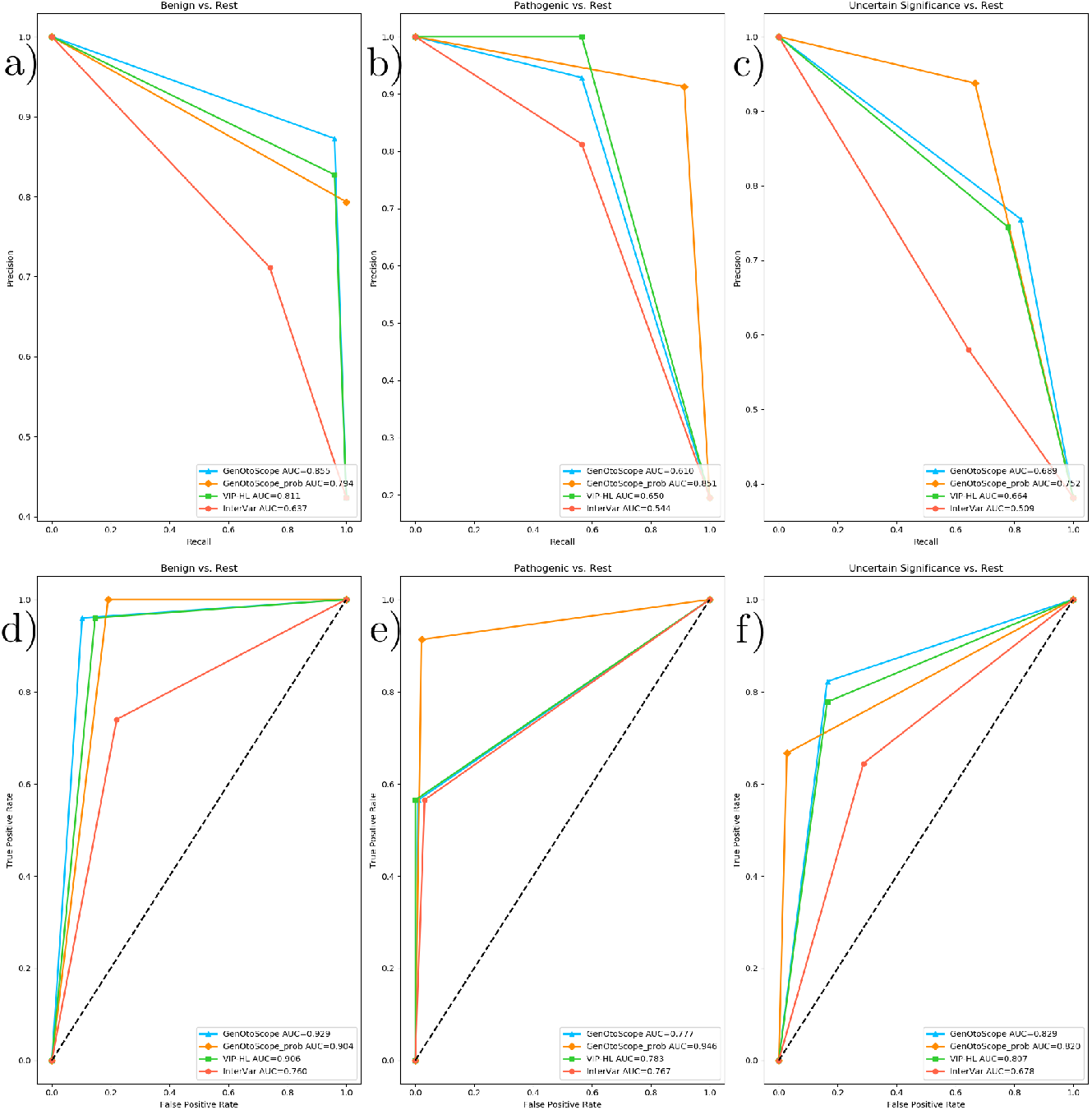
ROC curves and precision-recall curves for the MHH data set: a-c) ROC curve and AUC of all classification tools for VCEP-HL data set. a) Prediction of the “Benign” broader class versus the “Pathogenic” broader class and the VUS class b) Prediction of the “Pathogenic” broader class versus “Benign” broader class and the VUS class c) Prediction of the “VUS” class versus the “Benign” broader class and the “Pathogenic” broader class. d-f) Precision-recall curve and AUC of all classification tools for the MHH data set. d) Prediction of “Benign” broader class versus the “Pathogenic” broader class and the VUS class e) Prediction of the “Pathogenic” broader class versus “Benign” broader class and the VUS class f) Prediction of the “VUS” class versus the “Benign” broader class and the “Pathogenic” broader class.

We calculated the micro-average AUC of ROC curves and average precision of Precision-Recall curves, across the three broader classes for each classification tool. We show the results in Table 2. As in the previous data set, the two versions of the `GenOtoScope` classification achieved the best scores for both performance metrics.

**Table 2.**
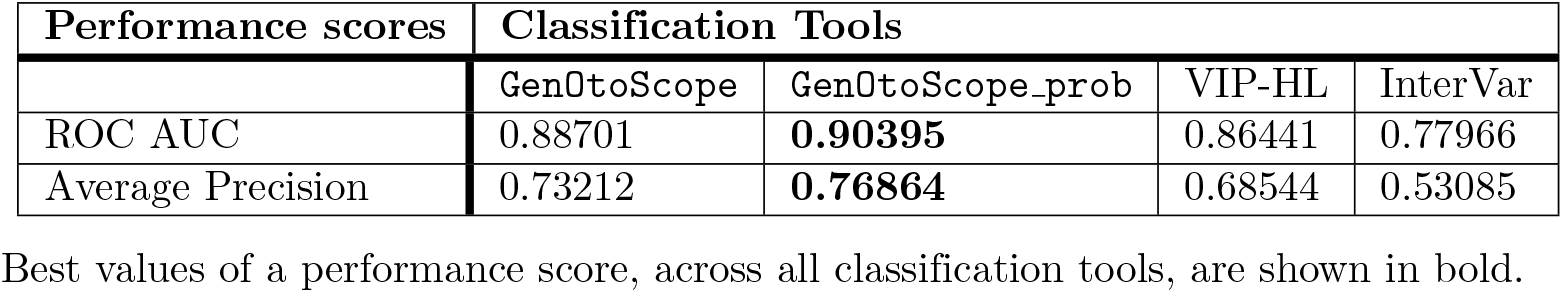
Micro-averaged performance scores for all classification tools, over the three broader classes in the MHH data set.

To explain the discrepancy in performance scores, we plotted the heatmap of log ratio of the activation frequency of a given tool compared to the activation frequency of the manual curation in Fig 9.

**Fig 9.**
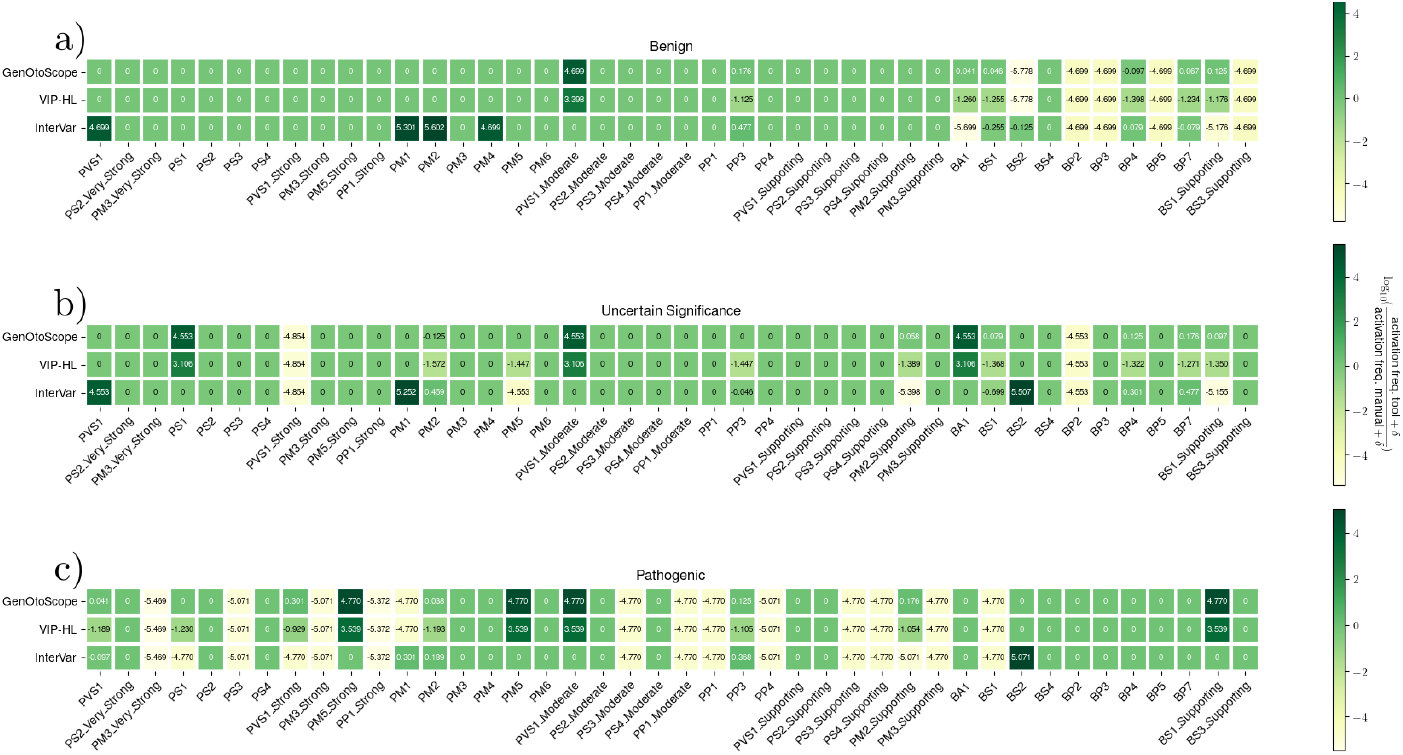
Activation frequency ratios for MHH data set. Log ratios calculated for each of the three classes classified by the MHH manual curators: (a) the “Benign” broader class, (b) the “VUS” class and (c) the “Pathogenic” broader class.

**Fig 10.**
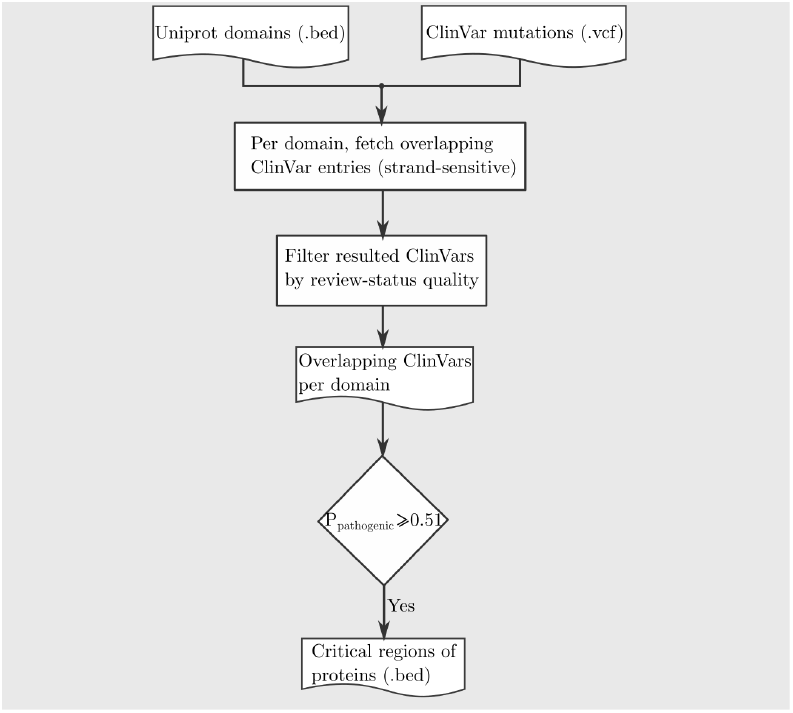
Conceptual workflow to call critical regions of proteins for assessment of PVS1 rule.

**Fig 11.**
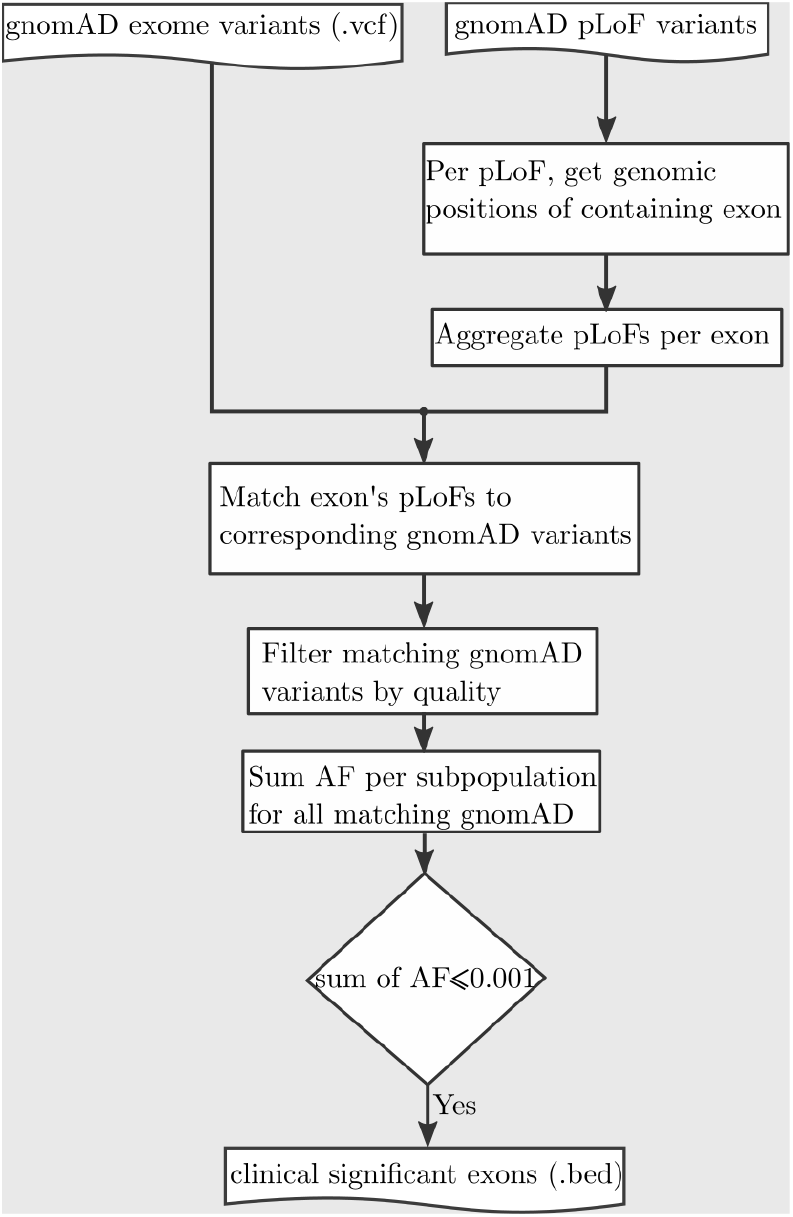
Conceptual workflow to call clinical significant exons for PVS1 rule.

For the “Pathogenic” broader class, VIP-HL activated 5 evidence-based rules (PVS1 (Strong), PP3, PM2, PS1 and PVS1) from 8 times less (PVS1 (Strong)) to 16 times less (PVS1). In contrast, for the same class, `GenOtoScope` activated 3 out of these 8 rules with the same frequency (PVS1, PS1 and PM2) as the manual curation. The remaining two rules were activated approximately one time more (PP3) and twice more often (PVS1 (Strong)) as the manual curation, respectively.

VIP-HL activated 8 implemented rules (BP7, BP4, BS1 (Supporting), BS1, PM2 (Supporting), PP3, PM5 and PM2) from 20 times less (BP4 and BP7) to 40 times less (PM2) than the manual curation for the “VUS” class. `GenOtoScope` activated 2 of these 8 rules (PM5 and PP3) with equal frequency to the manual curation.

`GenOtoScope` activated the remaining six rules (PM2, PM2 (Supporting), BS1, BP4, BS1 (Supporting) and BP7) with approximately one time more (BP7), up to one time less (PM2) as the manual curation.

For the “Benign” broader class, VIP-HL activated 6 rules (PP3, BS1 (Supporting), BP7, BS1, BA1 and BP4) from 12 times less (PP3) to 25 times less (BP4) than the manual curation. Contrary to VIP-HL pattern, `GenOtoScope` activated 2 out of these 6 rules (BA1 and BS1) with the same frequency as the manual classification and the remaining 4 rules (BP4, BS1 (Supporting), BP7 and PP3) with approximately two times more (PP3) up to one time less (BP4) as the manual curation.

Based on the observed motives on the activation frequency of each tool compared to the manual curation, we conclude that VIP-HL activated the aforementioned evidence-based rules less frequently than the manual curation. However, `GenOtoScope` was able to trigger the selected rules with similar or at most twice higher frequency compared to the manual curation. Consequently, we justify the best performance achieved in ROC and Precision-Recall scores by `GenOtoScope` for all three broader classes compared to the other two classification tools.

## Conclusion

In this work, we presented `GenOtoScope`, an automated classification tool for variants associated with congenital HL. Currently, our tool offers the classification through the automation of 12 out of 24 evidence-based criteria specified for HL [11]. We have shown that `GenOtoScope` outperformed other variant classification tools in terms of AUC score of ROC curve and of Precision-recall curve for all three broader classes (“Benign”, “VUS” and “Pathogenic”). To explain the difference in performance between the tools, we calculated the ratio of the activation frequency of triggered criteria by each tool and the manual curation. By comparing the ratios for each ACMG criterion, we observed that `GenOtoScope` achieved the most similar activation frequency to the manual curation, compared to the VIP-HL and InterVar tools.

Besides, the scope of this work is to provide an easily accessible tool to use for the classification of variants for HL phenotype. Therefore, we implemented two versions of the tool. The first version is a CLI application to be used by experienced bioinformatics personnel, who aim to classify a set of patients WES VCF files. Complementary, we have implemented a web interface to be used, by life scientists without bioinformatics expertise, to classify a single variant of interest. We hope that this tool will be used in search settings of genetic diagnostics routine diagnostic settings to provide a time-efficient and standardized classification of HL variants.

For future extension of `GenOtoScope` we aim to implement the most frequently activated evidence-based rules by manual curation to predict the two complementary grouped classes. For “Benign” broader class, the not implemented rules with highest activation frequency, by the manual curation, were the BS2, BP2, BP3, BP5 and BS3 (Supporting). For the “Pathogenic” broader class, the most frequently activated rules, only by the manual curation, were PM3, aggregated for all strengths, PP1, aggregated for all strengths, PS3 and PS4. To implement these rules, which need manual curation heavily, we aim to utilize database for genotype to phenotype such as DisGeNET [28] and prediction algorithms to link a mutation of interest to its respective functional study publications, for example AVADA [29]. Also, to facilitate even more no bioinformatics personnel to use the web interface, we would allow the user to input a single variant information on the welcome page without the need of creating a VCF file.

Last, by making the `GenOtoScope` an open source project, we aim to facilitate researchers to use its source code as a base to implement the ACMG evidence-criteria for phenotypes with a similar set of used evidence-based criteria, for example cardiomyopathy [4] or monogenic diabetes^2^.

## Supporting information

### S1 Appendix

#### PM1

The precise regions, used for PM1 criterion, are the pore-forming domain of KCN☯4 gene and the three-stranded helices of the collagen genes COL11A2, COL4A3, COL4A4 and COL4A5. PM1 is applied to missense variants overlapping any of the annotated genomic regions. If the variant overlaps on the three-stranded motifs of the collagen genes, it accepts only the matches that affect the Glycine residues contained in a Gly-X-Y motif.

#### PP3, BP4 and BP7

The used thresholds by prediction follow. To decide upon pathogenicity, we aggregated CADD and REVEL in the following scheme: if CADD score is greater than 20, then we set CADD_vote_ = 1 otherwise CADD_vote_ = 0. For REVEL, if REVEL score is greater or equal to 0.7, then REVEL_vote_ = 1, else if REVEL score is lower or equal to 0.15, then REVEL_vote_ = 0, otherwise we set REVEL_vote_ = 0.5. Finally, if the average voting of CADD_vote_ and REVEL_vote_ is greater or equal to 1, `GenOtoScope` assumes that the variant is pathogenic. For splicing impact, we aggregate the predictors MaxEntScan and dbscSNV in the following scheme: if 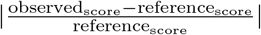 is greater than 0.15, then MaxEntScan_vote_ = 1 otherwise MaxEntScan_vote_ = 0. For dbscSNV, if either ADA score or RF score is greater than 0.6 then dbscSNV_vote_ = 1, otherwise dbscSNV_vote_ = 0. We aggregate the votes similarly to pathogenicity. That is, if the average voting of MaxEntScan_vote_ and dbscSNV_vote_ is greater or equal to 1, `GenOtoScope` decides that the variant has a splicing impact. Last for conservation prediction, we used PhyloP score, as follows: if PhyloP score is greater of 1.6 then `GenOtoScope` decides that this is a conserved site, otherwise `GenOtoScope` decides that the site is not a conserved site.

#### PM2 (PM2 Supporting), BA1 and BS1 (BS1 Supporting)

Regarding different inheritance patterns, the algorithm by default utilizes distinct thresholds for autosomal dominant and autosomal recessive inheritance mode as specified by [11]. For the X-linked mode of inheritance, autosomal dominant thresholds are adopted. If no mode of inheritance is provided, it is assumed to be unknown. In these cases, the algorithm selects the strictest threshold between autosomal dominant and recessive for each criterion. For mitochondrial genes, the same procedure is used as for unknown mode of inheritance, with an additional warning, since the application of ACMG criteria is validated only for Mendelian disorders.

### S2 Appendix

#### Constructing annotations for ACMG criteria

In the following, we explain how `GenOtoScope` utility scripts create the needed annotations for the automatic examination of the ACMG criteria. First, we present our methods to construct the annotations for PVS1 criterion: (i) critical regions for protein function, (ii) clinical significant exons and (iii) HL-relevant transcripts. Finally, we present how the respective `GenOtoScope` utility script creates the critical regions for protein function with no benign mutation.

#### Critical regions for protein function

To automate the assessment of PVS1 rule, the GenOtoScope’s sub-process construct three annotation files. In the first annotation file we include all critical regions for protein function. To create this file, the GenOtoScope’s respective sub-process maps all available ClinVar entries to the genomic positions of each UniProt domain, respecting the genomic strand of the domain. Then it filters-in all mapped ClinVar entries with at least 2 quality stars on their review status field. For each domain, it uses the interpretation field of the filtered-in overlapping ClinVar entries to calculate the probability that the region is pathogenic (and so critical for protein function) as:

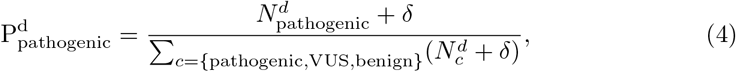

where 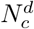 is the number of filtered ClinVar entries, found in the protein domain with UniProt id *d*, with class equal to *c* and *δ* = 10^−6^ is used as a smoothing parameter for the probability computation. Finally, the sub-process calls domains with 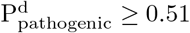 as critical for protein function. It saves all these critical domains for protein function in a BED file containing as columns: their genomic position, strand, their protein UniProt id and their corresponding 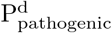 probability. The described procedure is also depicted in the Supplementary Fig 10.

#### Clinically significant exons

We also developed a sub-process to create an annotation file for clinically significant exons, which are exons at which loss of function variants are not frequent in the general population [13]. To do so, the sub-process first aggregates putative loss of functions (pLoF) variants of gnomAD [20] per Ensembl exon [21]. Second, for a given exon, it extracts the AF for each subpopulation of a pLoF variant intersecting the exon. Finally, it aggregates the AF of each extracted pLoF variant, for each subpopulation, and if this sum is lower than 0.001 for any subpopulation, the exon is called clinically significant. The output annotation file is in BED format containing the columns: the genomic position, strand, its exon Ensembl id and all containing transcript Ensembl ids, for each called clinical significant exon. The procedure is depicted in Supplementary Fig 11.

#### Hearing loss-relevant transcripts

The last annotation file for PVS1 is the hearing loss relevant transcripts. The respective sub-process utilizes three independent annotation files to create the hearing loss relevant transcripts and exons. The first file contains the phenotype relevant transcripts and their clinically relevant exons from [23], the second file contains disease-gene pairs for hearing loss from ClinGen repository [24] and the last file contains the clinical diagnostics panel for hearing loss created by the HG department of the MHH.

To aggregate these files, we argue that the [23] work contains the most detailed information as it contains not only hearing loss relevant transcripts for a given gene but also the clinically relevant exons of the respective transcripts. In contrast, the two remaining files contain either disease-gene pairs or only relevant transcripts of genes without specifying their clinically relevant exons. Therefore, the sub-process extends the relevant transcripts and clinically relevant exons reported by [23] with the relevant transcripts and all their contained exons from the diagnostic panel of HG department of MHH. Further, it extracts the longest coding transcript or the clinical relevant transcript annotated by Locus Reference Genomic resource (LRG) [25], and their contained transcripts, for all genes annotated by ClinGen but not found in the extended list of annotations of the last intermediate step. As final step, it aggregates the ClinGen unique genes annotations with [23] and the HG department of the MHH annotations to create the final hearing loss relevant transcripts and clinically relevant exons. The output annotation file is in BED format, containing the columns: chromosomal position, strand, transcript and exon ids for each clinically relevant exon.

#### Critical regions for protein function with no benign mutation

To automate PM1 evidence-based rule we need an annotation file containing the critical regions for protein function without benign mutations. To construct these annotated regions, we implemented a similar sub-process as for the critical regions used for PVS1 rule. The only difference is that this sub-process constraints the candidate domains with P_pathogenic_ *≤* 0.51 (Eq.4) to contain no benign ClinVar mutations. The resulting file is in BED format, containing the same columns as described above for the critical regions for protein function.

### Results

#### PVS1 annotations

`GenOtoScope` sub-processes created the three annotation files needed for the refined PVS1 criterion. For the first file, critical regions, we applied our methodology using the 25,552 ClinVar entries, version of March 2021 and 12,776 UniProt domains, version of February 2021. The resulting file contains 1,478 UniProt domains annotated as critical for protein function. Using the HL-relevant transcripts and exons curated in [23], [24] (VCEP-HL) and MHH diagnostic panel, we extracted 2,812 unique exons and 215 unique transcripts, contained in 154 genes. For the annotation file with the clinical significant exons, the used version of the pLoF variants and the allele frequency of exomes of gnomAD was the version 2.1.1. By this process, we annotated 107,966 exons as clinical significant exons.

#### PM1 annotations

The 10 mutational hotspots relevant to HL as published by VCEP-HL committee, at page^3^, were utilized to create the annotation file to evaluate the PM1 criterion. Using the ClinVar entries and UniProt domains, same versions as described above. Besides 750 UniProt domains were called as critical regions for protein function without a benign variant. To evaluate the PM1 evidence-criteria we intersect the chromosomal position of the input variant with both annotation files.

### S3 Appendix

#### Implementation note

To automate the examination of the ACMG evidence-based criteria we used the Python programming language. First, we used the hgvs library [30] to parse the variant information for each Ensembl transcript. Then, using the HGVS format we constructed the observed coding sequence and we extracted the needed information for the criteria, through the PyEnsembl library^4^. For example, we extracted the start and stop positions of exons and the positions of the start and stop codons. This library uses the Ensembl version 75 for GRCh37 human genome. We used the PyVCF library^5^ to parse variants from VCF files, for instance the ClinVar variants. We applied the Pybedtools [31] to find the intersections of annotation files, such as the overlap of UniProt domains with repeat regions. Finally, we utilized the BioPython library [32] for all other tasks for example, to convert cDNA codons to amino acids. `GenOtoScope` currently works only for grch37 genome assembly coordinates. The performance metrics were calculated using scikit-learn library [40].

The bioinformatics user shall download the whole tool code along with the set of data needed for its execution, e.g. annotation files HL or known variants with high MAF from the github repository. Besides, at this repository, the user can find example configuration files, example input with the corresponding output files and a documentation on how to install and execute `GenOtoScope` on a linux machine or server. Last, to be able to use the variant annotation script, genotoscope annotate.py, the user needs to install the megSAP application on a docker container, as explained on the respective tool github repository ^6^.

## Disclaimer

The classification produced by `GenOtoScope` is intended for an efficient pathogenicity prediction of WES files, thus for research use only. It is not intended for diagnostic or clinical purposes. The classification provided by `GenOtoScope` does not replace a physician’s medical judgment and usage is entirely at your own risk. The providers of this resource shall in no event be liable for any direct, indirect, incidental, consequential, or exemplary damages.

## Acknowledgements

The authors would like to acknowledgement the financial support through the *Understanding Cochlear Implant Outcome Variability using Big Data and Machine Learning Approaches*, project id: ZN3429, funded by the Volkswagen Foundation, through the Ministry for Science and Culture of Lower Saxony Germany (MWK: Ministerium für Wissenschaft und Kultur). DPM would like to express many thanks to Oleh Astapiev, Christos Mauromatis and Sotirios Mauromatis for help on setting up the web interface. Equally, DPM would like to thank Anna-Lena Katzke and Dr. Winfried Hofmann for installing the `GenOtoScope` web interface in the MHH server system.

## Author Contributions

Conceived the idea: BA. Designed the experiments: DPM CL GS ASH. Recruited the HL patients cohort (MHH data set): ALS SvH CL. Classified variants of HL patients cohort (MHH data set): ASH SvH CL GS DA. Performed the experiments: DPM. Wrote the paper: DPM. Critically revised the manuscript CL GS ASH SvH BA WN ALS. Developed the algorithms and the software: DPM. Tested the algorithms and the software: DPM CL GS ASH.

## Footnotes

1 https://github.com/imgag/megSAP

2 https://clinicalgenome.org/site/assets/files/7039/clingen_diabetes_acmg_specifications_v1.pdf

3 https://submit.ncbi.nlm.nih.gov/ft/byid/vroiax8b/hearing_loss_acmg_specifications_v1_2018.pdf

4 https://github.com/openvax/pyensembl

5 https://github.com/jamescasbon/PyVCF

6 url https://github.com/imgag/megSAP

